# EVALUATION OF ANTI-BACTERIAL AND ANTHELMINTIC ACTIVITIES OF BARK OF PLANT *CASSIA FISTULA* Linn

**DOI:** 10.1101/2020.10.19.345520

**Authors:** Anant Nayabaniya, Samyam Aryal, Bibek Dahal, Hemanta Khanal, Anil K. Gupta

## Abstract

This study deals with phytochemical screening and evaluation of anti-bacterial and anthelmintic activities of the bark extracts of *Cassia fistula* L. The extraction with different solvents - methanol, acetone, distilled water - had been carried out. The anti-bacterial assay was done against two gram +ve and two gram −ve bacteria by agar cup diffusion method. The anthelmintic activity was done on *Pheretima posthuma* (earthworm) by recording the paralyzing and death time under different concentrations. Methanolic extract showed the maximum activity on both fronts be it anti-bacterial or anthelmintic. The extract was effective only against gram +ve bacteria. Among *Staphylococcus aureus* and *Streptococci faecalis*, the former was more susceptible. The activity against gram −ve bacteria was not found. In case of anthelmintic activity, the concentration of 50 mg/ml was effective and near to the paralyzing and death time as recorded for the 10 mg/ml concentration of the standard Albendazole. However, aqueous extract was more effective than the acetone extract.

## 1. Introduction

Herbalism and ethnobotany has a growing impact on the present biomedical research. There has been a shift in paradigm as natural products find an increased entry into clinical trials [1]. Natural products are also sought in the drug development phase as potential leads as anti-neoplastic and anti-microbial agents [2]. The shift could lead to the advancement of traditional and ethnical medicine for an enquiry through a scientific-lens not only leading to a safer, rational, and effective usage but also towards preserving the traditional heritage and generation of income [3].

Estimates show that more than 25% of the conventional medicines are derived from plant sources [4]. This combined with the fact that nearly 80% of the developing and upwards 50% of the developed world relies on natural products warrants a more intrigued and scientific attitude towards herbaceuticals[5][6].

*Cassia fistula* Linn., commonly known as Amulthus or “Indian Laburnum” is a plant, whose derivatives that that is frequently prescribed in Ayurveda – the traditional medical system of the Indian subcontinent[7]. Traditionally the root and the fruit pulp have been used as a form of stimulant laxative and as an anti-helminthic. The roots of the plant are also prescribed as a tonic, and an anti-pyretic [8]. The extract of the pods is known to reduce blood sugar in animal models [9]. The leaves are externally applied for dermatophytosis and eczema. The fruits are reported to be used for asthma. The seeds possess emetic and cathartic properties while the powder of seeds traditionally was being used as anti-amebic agent [10][11].

Owing to these claims various studies have been conducted to explore the pharmacological properties of the plant. The alcoholic extract of the stem bark showed inhibition of *S. aureus* compared to aqueous extract [12]. The aqueous and ethanolic extracts could potentially be used as anti-microbial agent for various infections. Methanolic extract of of *Cassia fistula* L. assayed for in vitro antibacterial activity against eleven pathogenic bacteria by using the diffusion method in agar showed effects equivalent to standard antibiotics [13]. Also, the extracts from seeds and the pulp at the concentration of 100mg/ml showed significant paralysis and death of worms (*Pheretima posthuma*) as compared to reference drug Piperazine citrate[14].

A rodent study on ethanolic extracts of *Cassia fistula* L. at various doses (50, 100, 250, 500 and 750 mg/kg by weight) showed anti-inflammatory effect which was compared with standard anti-inflammatory drugs (Diclofenac and Indomethacin). The ethanolic extracts showed observable anti-inflammatory as well as anti-pyretic effect [15].

Previous studies have shown significant anti-micorbial and anti-pyretic effect of various parts of *Cassia fistula*. However, the bark of the plant remains significantly understudied. As such this study aims to delineate the anti-bacterial and anti-helminthic properties of *Cassia fistula*.

## 2. Materials and Methods

### 2.1 Plant collection and authentication

The bark of plant Cassia fistula was collected locally from the forest of Chinde danda, Dharan-24, Sunsari, Nepal during the month of July to August 2016 and authenticated by Department of Botany, Post Graduate Campus, Biratnagar, Nepal. The voucher specimens were deposited in the same Department. (Ref no. 073/074-12)

### 2.2 Extraction

First, the plant materials were cut into small sections and were shade dried at room temperature and then were dried in a hot air oven 65°C for 10 hours. The barks were intermittently inspected to appreciate if the drying was complete. The dried sample was then crushed into powder by electric grinder and the coarse powder was passed through a 40-mesh sieve to collect the fine powder which was subjected to extraction by using suitable solvents in the soxhlet apparatus followed by evaporation of solvent and drying the extract and storing at 4° C. 25gm dried powder of the stem bark was extracted with 160ml of each solvent – methanol, acetone, and distilled water. The concentrated products were weighed and the yield was calculated. Then the extract was passed through Whatman No.1 filter paper and the filtrate was concentrated by evaporation in order to reduce the volume. The paste like extract was stored in labeled screw capped bottles and kept in refrigerator at 4°C.

### 2.3 Preliminary phytochemical Analysis

The dried barks of *Casssia fistula L.*, was grind to powder using an electric blender and dissolved separately in 100 ml of solvent. This solution was kept under room temperature for seven days to allow the extraction of compounds from the stem bark. The solution of each sample was stirred after every 24 hours using sterile glass rods. After seven days, the solution was filtered through what man No-1filter paper. The solvent was evaporated and sticky substance obtained that was stored in refrigerator and suspended in 10% (DMSO) Di-methyl Sulfoxide prior to use. Chemical tests were carried out with both the plants extracts and on the powder specimens using standard procedure to identify the constitutions as described by Harborne, the specific procedure involved for the evaluations of a particular group of chemical is mention below.

#### 2.3.1 Tannins

1 ml of water and 1-2 drops of ferric chloride solution were added in 0.5ml of extracted solution. Blue color was observed for tannis and green black for methanolic tannin.

#### 2.3.2 Saponins (Foam test)

Small amount of extract was shaken well with little quantity of water. If foam produced persist for ten minutes if the presence of saponins.

#### 2.3.3 Flavonoids (Alkaline Reagent test)

Extractions were treated with few drops of sodium hydroxide solution. Formation of intense yellow color, which because colors on addition of acid, Indicates the presence of flavonoids.

#### 2.3.4 Steroids

2 ml of acetic anhydride was added to 0.5g extract of each sample with 2 ml H_2_SO_4_. The color changed from violet or blue or Green in some samples indicating the presence of steroids.

#### 2.3.5 Terpenoides (salkowski test)

5 ml of each extract was mixed in 2 ml of chloroform, and concentrated H2SO4 (3ml) was carefully added to form a layer. A reddish brown coloration of the interface was formed to show the presence of terpenoids.

#### 2.3.6 Cardiac glycosides (Keller-Killani test)

5 ml of each extract was treated with 2 ml of glacial acetic containing drop of ferric chloride solution. This was underplayed with 1 ml of concentrated sulphuric acid. A brown ring of inter face indicates a deoxysugar characteristics of carotenoides. A violet ring may appear below ring while in the acetic layer, a greenish ring may from just gradually throughout thin layer.

#### 2.3.7 Alkaloids

Alkaloids are basic nitrogenous compounds with definite physiological and pharmacological activity. Alkaloids solution produces while yellowish precipitated a few drops of Mayer’s reagents are added.

#### 2.3.8 Anthraquinones

Born trigger’s test will be used for detecting the presence of anthroquinone. In this case 0.5g of planet extract was shaken with benzene layer separated and half of its own volume of 10% ammonia solution added. A pink, red, or violet coloration in the ammonia phase indicated the presence of anthraquinone.

### 2.4 Test Organisms

The pathogenic bacterial species (S. *aureus*, S. *faecalis*, E. *coli*, Salmonella *typhi*) were collected from the Department of Microbiology, B.P. Koirala Institute of Health Sciences (BPKIHS), Dharan, Sunsari, Nepal. The adult earthworms *Pheritima posthuma* were made acquired from the Laboratory of Microbiology of Sunsari Technical College, Dharan, Sunsari and washed with normal saline to remove all fecal matter and dirt.

### 2.5 Anti-microbial assay

Anti-microbial activity was tested using agar cup diffusion method ^[16]^. The petridish were first prepared with 20ml Muller Hinton agar (Hi-media, Mumbai). The test cultures were inoculated on top of solidified media using an inoculation loop. The cups were prepared with the help of a borer (6mm). They were then filled with solution of suitable concentration of sample (800 mcg/ml, 400 mcg/ml, 200 mcg/ml, 100 mcg/ml, 50 mcg/ml, 25 mcg/ml – 1 ml each for each solvent extracts). Chloramphenicol (30mcg/disc) was used as positive control. The plates were incubated at 37 o C for 24 hours. The zone of Zone of inhibition for each test and control were recorded in millimeters

### 2.6 Anti-helminthic activity

The earthworms 3-5 cm in length and 0.1-0.2 cm in width were used for the experimental protocol. A differing concentration of plant extracts were prepared in distilled water. The concentrations were 10 mg/ml, 20 mg/ml, 25 mg/ml and 50 mg/ml for each solvent extracts. The standard drug, Albendazole, was prepared in distilled water at a dose level of 10 mg/ml. The earthworm serving as negative control received distilled water. The earthworms were kept in separate petri dish for each test and control samples and were treated with respective solutions. Observation was made for the time taken to cause paralysis and death of individual worms. Paralyzing and death time were concluded when the worms lost their motility followed with fading away of their body colours [14].

## 3. Result

Phytochemical screening of the plant showed the presence of different constituents in different solvent extracts. The phytochemical analysis showed that the various groups that were found to be present in the different extracts are listed in table 1.

**Table 1.** Result of phytochemical screening

| S.N. | Tests | Methanol extract | Acetone extract | Distilled water extract |
| --- | --- | --- | --- | --- |
| 1. | Tannin | + | + | + |
| 2. | Saponin(foam test) | + | + | + |
| 3. | Flavonoids(Alkaline reagent test) | + | + | + |
| 4. | Steroids | - | - | - |
| 5. | Terpenoids | + | + | + |
| 6. | Cardiac glycoside(keller killani test) | - | - | - |
| 7. | Alkaloids | - | - | - |
| 8. | Anthraquinones(Born trigger's test) | + | + | + |

The methanolic, acetate, and aqueous extract at higher concentration (>50mcg/ml) of *Cassia fistula* stem bark showed significant inhibitory zone against gram positive S. aureus and S. fecalis. The gram-negative bacteria were not inhibited (table2). These actions are in accord to previous such studies done in the plant.

**Table 2:**
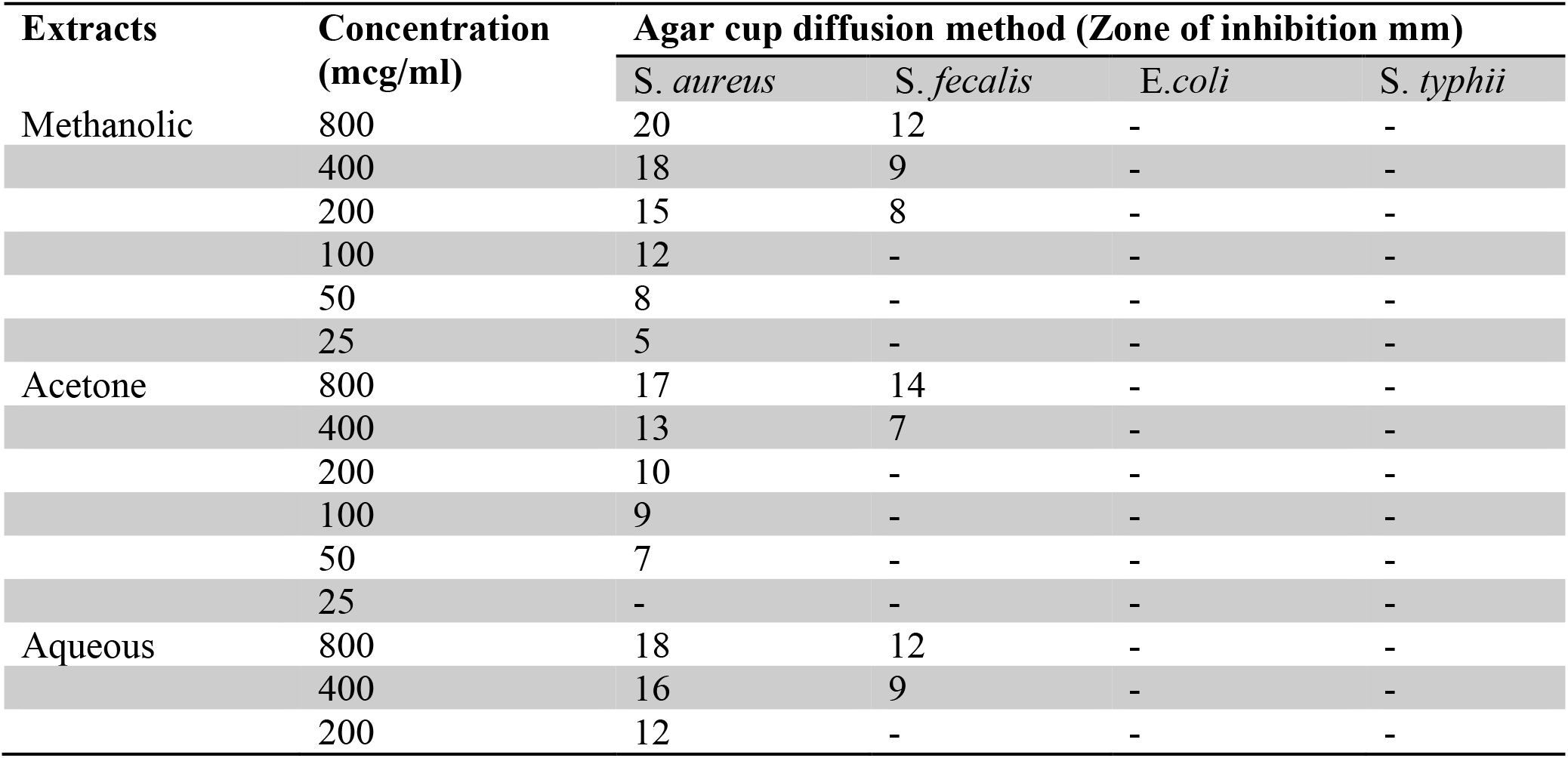
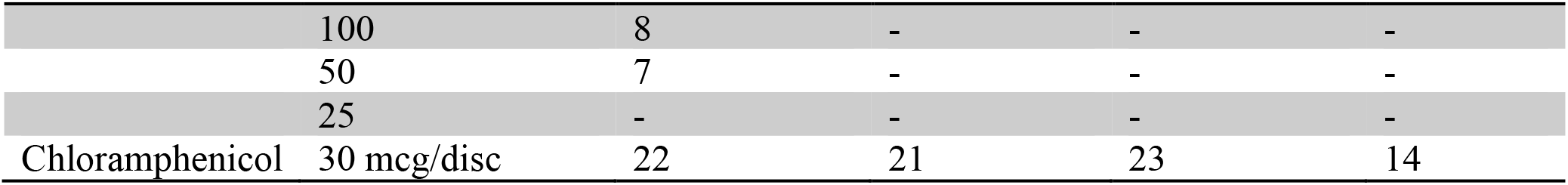
Antibacterial activity of various solvent extracts of *Cassia fistula* stem bark

Time to paralyze and death for individual worms, *Pheretima posthuma,* were observed. Time for paralysis was noted when no movement were observed except when the dish were shaken vigorously. When there was a loss of motility and fading of body color the worm was concluded to be dead (table3).

**Table 3:** Antihelminthic activity of various solvent extracts of *Cassia fistula* stem bark

| Extracts | Concentration (mg/ml) | Paralyzing time (minutes) | Death time (minutes) |
| --- | --- | --- | --- |
| **Methanolic** | 10 | 49 | 132 |
|  | 20 | 35 | 93 |
|  | 25 | 28 | 81 |
|  | 50 | 20 | 67 |
| **Acetone** | 10 | 58 | 139 |
|  | 20 | 41 | 122 |
|  | 25 | 37 | 97 |
|  | 50 | 25 | 77 |
| **Aqueous** | 10 | 55 | 135 |
|  | 20 | 39 | 119 |
|  | 25 | 33 | 93 |
|  | 50 | 24 | 79 |
| **Standard albendazole** | 10 mg/ml | 16 | 28 |
| **Blank 6% DMSO water** | - | - | - |

## 4. Discussion

The methanolic extract of the stem bark exhibits the maximum anti-bacterial activity. Previous study had also demonstrated that the methanolic extract of the bark is more efficient than aqueous extract against S. aureus [12]. Previous study had also demonstrated that the extracts of flowers are effective against B. *cereus*, S. *aureus*, S. *epidermidis*, E. *coli* and K. *pneumoniae*. The most susceptible microorganisms to methanolic extracts were E. *coli* and K. *pneumoniae* respectively. Also, On the contrary, S. *aureus* and B*. cerus* showed least sensitivity to methanolic extracts respectively [17]. This is the first study demonstrating the activity of various extracts of the stem bark against gram positive bacteria. Previous study on flower extract had shown activity against Gram positive bacteria and P. *aeruginosa* [18]. This aligns with the finding that gram negative bacteria are more resistant to plant products than gram positive bacteria [19]. This, also, demonstrates that the extract can possibly be used as a narrow spectrum antibiotic for *Staphylococcus aureus*.

The anthelmintic activity of C. *fistula* may be due to phenolic compound, tannins that can bind to glycoprotein on the cuticle of the parasite and possibly causing death [20]. In this study, the shortest time of paralysis and death was found at dose 50 mg/ml which was possibly comparable to the paralyzing and death time for albendazole. Previous study had reported shortest time of paralysis and death at dose of 100 mg/ml using fruit pulp and seed extracts which was near to the paralyzing and death time as recorded for the 10 mg/ml concentration of the standard piperazine citrate [14]. This is a first study comparing various stem bark extracts against the widely used vermicidal – albendazole.

The study has several limitations, since, serial broth dilution was not performed, the Minimum Inhibitory Concentration (MIC) for various bark extracts remains unclear. This becomes necessary specially during the use in clinical settings.

## Conclusion

The assessed properties of *Cassis fistula* stem bark extract by the present work, substantiates the various claims obtained from ethno botany and Ayurveda including the use in diarrhea, skin infections, and worm infestations. The plant can possibly be a useful narrow spectrum antibiotic against Staphylococcus aureus and can possibly be used as a vermicidal agent.

## 5. Conflict of interest

None

## Notes

### Competing Interest Statement

The authors have declared no competing interest.

